# The *Drosophila* mitochondrial citrate carrier regulates L-2-hydroxyglutarate accumulation by coupling the tricarboxylic acid cycle with glycolysis

**DOI:** 10.1101/297705

**Authors:** Hongde Li, Alexander J. Hurlburt, Jason M. Tennessen

## Abstract

The oncometabolites D- and L-2-hydroxyglutarate (2HG) broadly interfere with cellular metabolism, physiology, and gene expression. A key regulator of 2HG metabolism is the mitochondrial citrate carrier (CIC), which, when mutated, promotes excess D-/L-2HG accumulation. The mechanism by which CIC influences 2HG levels, however, remains unknown. Here we studied the *Drosophila* gene *scheggia* (*sea*), which encodes the fly CIC homolog, to explore the mechanisms linking mitochondrial citrate efflux to L-2HG metabolism. Our findings demonstrate that decreased *Drosophila* CIC activity results in elevated glucose catabolism and increased lactate production, thereby creating a metabolic environment that inhibits L-2HG degradation.

## Introduction

The enantiomers of 2-hydroxyglutarate (2HG) have emerged as potent regulators of metabolism, chromatin architecture, and cell fate decisions (Losman and Kaelin 2013; Ye et al. 2018). While these compounds are commonly associated with cancer metabolism and often referred to as oncometabolites, neither D-2HG nor L-2HG are tumor specific. Humans produce D-2HG as the result of γ-hydroxybutyrate metabolism and phosphoglycerate dehydrogenase activity (Struys et al. 2005b; Fan et al. 2015), while L-2HG is generated by malate dehydrogenase and lactate dehydrogenase A (LDHA) in response to hypoxia, acidic cellular conditions, and decreased electron transport chain (ETC) activity (Mullen et al. 2011; Reinecke et al. 2012; Intlekofer et al. 2015; Oldham et al. 2015; Nadtochiy et al. 2016; Teng et al. 2016; Intlekofer et al. 2017). Furthermore, both yeast and *Drosophila* generate D-2HG and L-2HG under standard growth conditions, respectively (Becker-Kettern et al. 2016; Li et al. 2017), suggesting that these molecules serve endogenous biological functions and emphasizing the need to understand how 2HG metabolism is controlled *in vivo*.

Despite the fact that D- and L-2HG dramatically influence cellular physiology, the molecular mechanisms that regulate 2HG accumulation in healthy cells remain poorly understood. In fact, most of our current understanding about endogenous D- and L-2HG metabolism stems from a class of rare human diseases that are collectively known as the 2HG acidurias (2HGAs, Kranendijk et al. 2012). For example, patients with L2HGA accumulate L-2HG due to loss-of-function mutations in the FAD^+^-dependent enzyme L-2HG dehydrogenase (L2HGDH) (Rzem et al. 2004), which converts L-2HG to 2-oxoglutarate (2OG). Similarly, D2HGA type I results from the absence of D-2HG dehydrogenase (D2HGDH) activity and an inability to degrade D-2HG (Struys et al. 2005a). Overall, these studies illustrate how the 2HGA disorders provide essential clues for understanding endogenous 2HG metabolism.

In addition to the disorders associated with a single 2HG enantiomer, a small subset of 2HGA patients exhibit elevated levels of both D-2HG and L-2HG. This rare disease, which is known as combined D-/L-2HGA, results in severe neurological and muscular defects, developmental delays, and childhood lethality (Muntau et al. 2000). Intriguingly, combined D-/L-2HGA results from loss-of-function mutations in *SLC25A1* (Nota et al. 2013), which encodes a mitochondrial citrate carrier (CIC) that mediates the transport of citrate across the mitochondrial inner membrane (Palmieri 2013). Based on both the metabolic profile of combined D-/L-2HGA patients and stable-isotope tracer studies of CIC-deficient cells, this transporter coordinates the tricarboxylic acid (TCA) cycle flux with lipogenesis and cellular redox balance (Mullen et al. 2011; Jiang et al. 2017). The mechanism by which CIC influences D-2HG and L-2HG accumulation, however, remains largely unknown.

We recently discovered that *Drosophila* larvae accumulate high concentrations of L-2HG during normal larval growth (Li et al. 2017). Subsequent analysis revealed that larvae rely on the *Drosophila* LDH homolog (dLDH) to synthesize L-2HG from the TCA cycle intermediate 2-oxoglutarate (2OG) (Li et al. 2017). Considering that human LDHA also synthesizes L-2HG (Intlekofer et al. 2015; Nadtochiy et al. 2016; Teng et al. 2016; Intlekofer et al. 2017), our earlier study indicates that *Drosophila* is well suited to explore the basic metabolic mechanisms that control L-2HG accumulation. Here we exploit the fly system to investigate the metabolic link between CIC activity and L-2HG. By studying a hypomorphic mutation in the *Drosophila* gene *scheggia* (*sea*), which encodes the fly *SLC25A1* homolog (Carrisi et al. 2008; Morciano et al. 2009), we demonstrate that loss of mitochondrial citrate efflux results in elevated glucose catabolism, increased lactate production, and enhanced L-2HG accumulation. The dramatic increase in L-2HG levels, however, primarily result from decreased degradation, as the increase in lactate concentration inhibits dL2HGDH activity and stabilizes the L-2HG pool. Overall, our study reveals a mechanism by which a well-described metabolic feedback loop unexpectedly controls L-2HG degradation.

## Results and Discussion

### L-2HG levels are increased in sea mutants

To determine if the *Drosophila* homolog of *SLC25A1* influences D-2HG and L-2HG accumulation, we used gas chromatography-mass spectrometry (GC-MS) to quantify the 2HG enantiomers in *sea* mutant larvae (*sea*^∆24^/*Df*) and genetically matched heterozygous controls (*sea^Prec^/Df*). Our analysis revealed that both D- and L-2HG are significantly elevated in *sea* mutants (Fig. 1A,B), with L-2HG representing the majority of the 2HG pool within *sea*^∆24^/*Df* larvae. While these observations differ from combined D-/L-2HGA patients, where D-2HG is the more abundant enantiomer (Muntau et al. 2000), the metabolic profile of *sea^∆24^/Df* mutants clearly indicates that the inverse relationship between CIC activity and L-2HG accumulation is present in flies.

**Figure 1.**
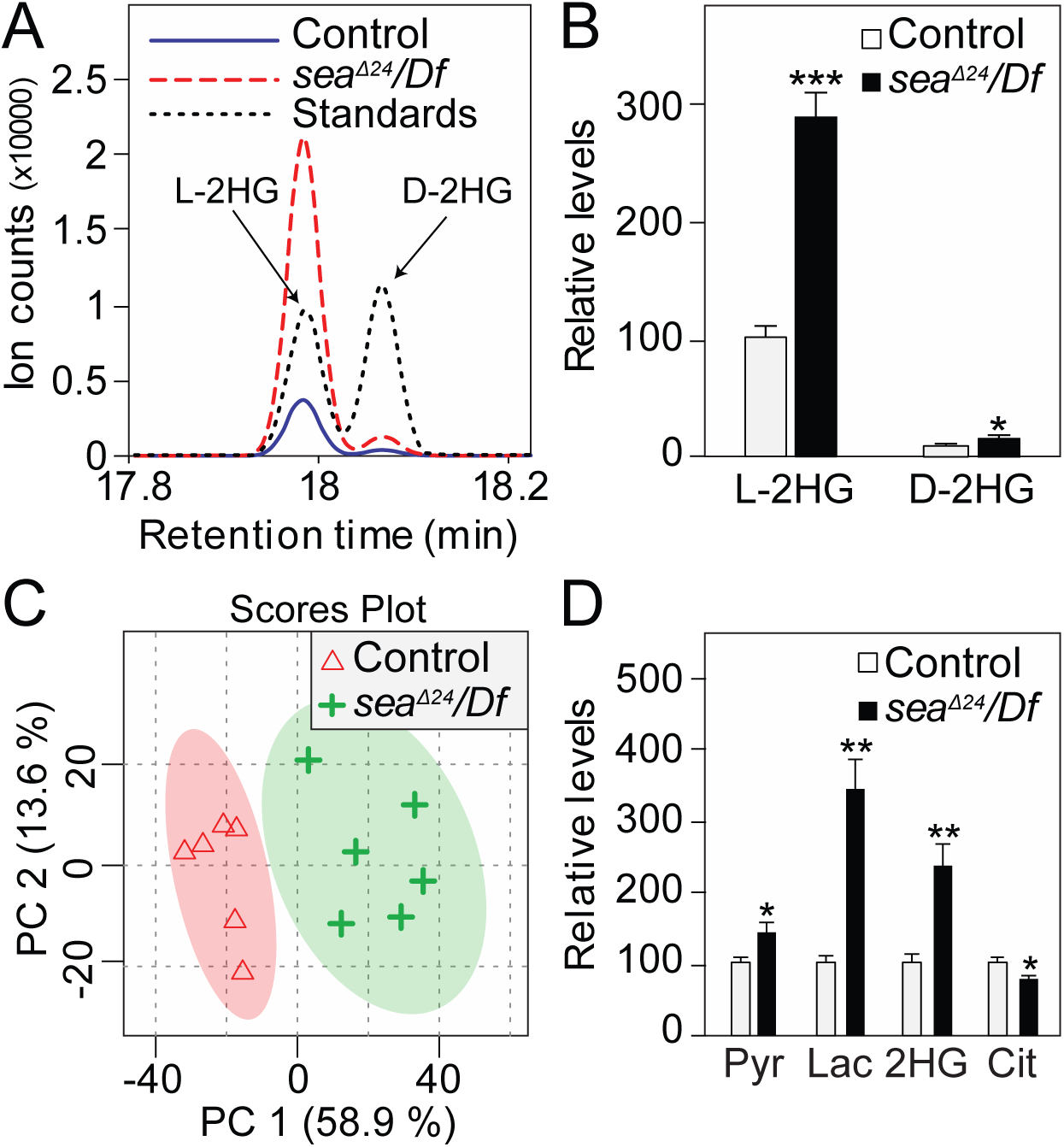
*sea* mutant larvae accumulate excess L-2HG. (**A**) L- and D-2HG in larvae were detected separately using a chiral derivatization method coupled with GC-MS. (**B**) Relative abundance of L-2HG and D-2HG in *sea* mutant and control larvae. (**C**) The PCA scores plots of GC-MS spectra show that the metabolic profile of *sea*^Δ24^/*Df* mutants is significantly different than that of the *sea*^prec^/*Df* control. (**D**) Targeted analysis of the GC-MS data analyzed in panel **(C)** reveals that *sea*^Δ24^/*Df* mutants display significant changes in pyruvate (pyr), lactate (lac), 2-hydroxyglutarate (2HG) and citrate (cit). For all panels, data are shown as mean ± SEM, *n* = 6, \**P* < 0.05, \*\**P* < 0.01.

#### sea mutants and combined D-/L-2HGA patients exhibit a similar metabolic profile

Combined D-/L-2HGA patients not only exhibit increased 2HG levels and decreased citrate accumulation, but also possess elevated levels of lactate, 2OG, succinate, fumarate, and malate (Nota et al. 2013; Prasun et al. 2015). To determine if *sea* mutants exhibit similar metabolic defects, we used a GC-MS-based approach to examine metabolites in glycolysis and the TCA cycle. Multivariate data analysis of the resulting datasets revealed that *sea*^∆24^/*Df* mutant larvae exhibit a distinct metabolic profile when compared to either genetically-matched *sea*^Prec^/*Df* controls or *w*^1118^/*Df* controls (Fig. 1C; Supplemental Fig. S1A), demonstrating that the *sea*^∆24^ mutation significantly disrupts larval metabolism. Targeted analysis of these data revealed that *sea*^∆24^/*Df* mutants display a metabolic profile that is reminiscent of combined D-/L-2HGA patients. Notably, mutant larvae exhibit decreased citrate levels and elevated amounts of pyruvate, lactate, fumarate, and malate (Fig. 1D; Supplemental Fig. S1B). Similar metabolic changes were observed when the *sea*^∆24^ mutation was analyzed in trans to a second deficiency that also uncovers the *sea* locus (Supplemental Fig. S2). Moreover, the *sea*^∆24^/*Df* metabolic phenotypes were rescued by ubiquitous expression of a *UAS-sea* transgene, indicating that the metabolic profile displayed by *sea*^∆24^/*Df* mutants specifically results from the loss of CIC activity (Supplemental Fig. S3A,B).

#### sea mutants exhibit elevated glycolytic flux

Considering that *Drosophila* larvae primarily synthesize L-2HG from glucose (Li et al. 2017), our data suggest that the *sea*^∆24^/*Df* metabolic profile results from elevated glycolytic flux. We tested this hypothesis by feeding ^13^C_6_-glucose to both *sea*^∆24^/*Df* mutants and *sea*^prec^/*Df* controls, and selectively monitoring ^13^C incorporation into lactate, pyruvate, citrate, and 2HG. When compared with *sea*^Prec^/*Df* controls, *sea*^∆24^/*Df* mutants exhibit a 60% increase in the rate of lactate (m+3) synthesis. Moreover, mutant larvae also exhibited a modest increase in the production of pyruvate (m+3) and 2HG (m+2), and a slight, but significant, decrease in the rate of m+2 citrate synthesis (Fig. 2). These observations confirm that glycolytic flux is elevated in *sea* mutants and are consistent with a recent study that observed enhanced glucose consumption and increased lactate production in CIC-deficient human cells (Jiang et al. 2017).

**Figure 2.**
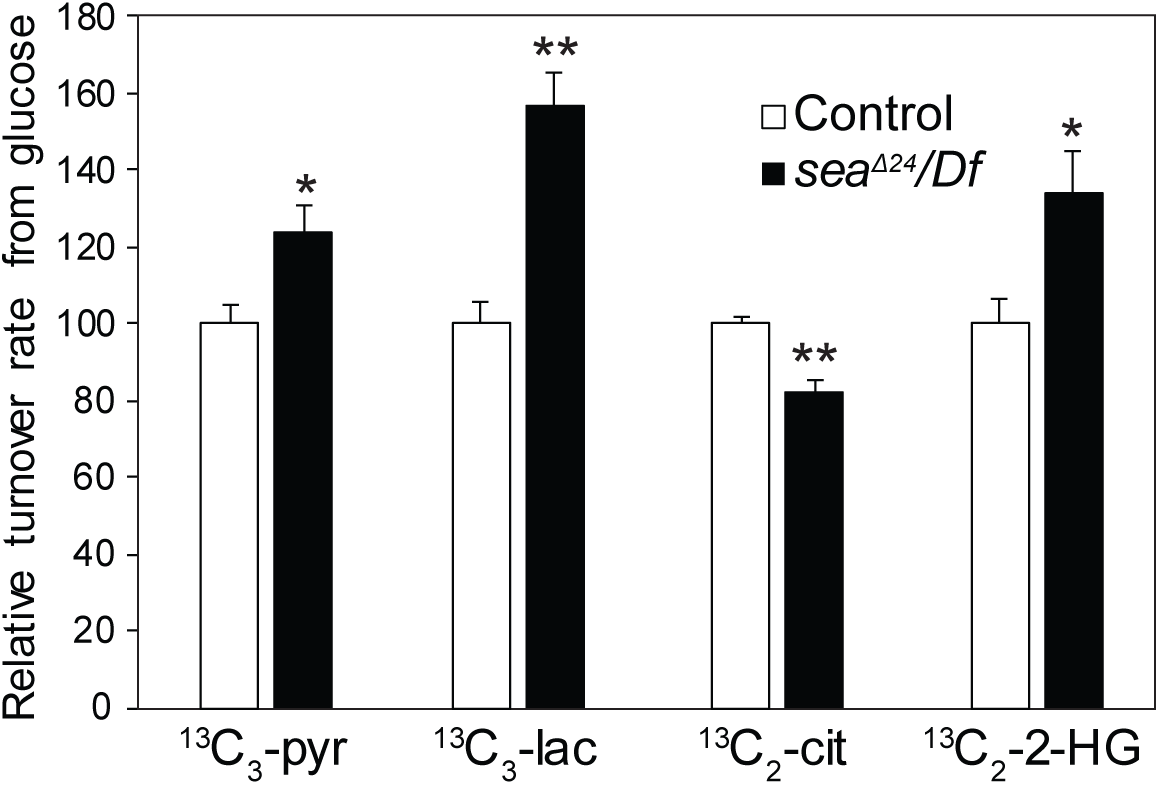
*sea* mutants exhibit elevated levels of glycolytic flux. The relative metabolic flux rates from ^13^C_6_-glucose into pyruvate (pyr; m+3), lactate (lac; m+3), 2HG (m+2), and citrate (cit; m+2). Data are shown as mean ± SEM, *n* = 4, \**P* < 0.05, \*\**P* < 0.01.

The ^13^C tracer experiments raise the question of how *Drosophila* CIC activity antagonizes glucose catabolism and L-2HG accumulation. Since *sea* mutants exhibit significant changes in metabolites associated with histone modifications (*i.e*., cytosolic citrate and L-2HG), we used RNA sequencing (RNA-seq) to determine if *Drosophila* CIC activity regulates the expression of glycolytic enzymes. When compared to *sea*^Prec^/*Df* controls, *sea*^∆24^/*Df* mutants exhibited significant changes in ~1,800 genes (fold change >2.0; p-value <0.01; Supplemental Table S1). However, out of 25 genes that encode glycolytic enzymes, the mRNA levels for 23 were either unchanged or slightly reduced in *sea*^∆24^/*Df* mutants (Supplemental Table S2). Moreover, *hexokinase C* (*Hex-C*) and *triose phosphate isomerase* (*Tpi*) were the only glycolytic genes that exhibited a >1.5-fold increase in *sea*^∆24^/*Df* mutants (no gene in this 25-gene subset exhibited a greater than a 2-fold increase). Similarly, qRT-PCR confirmed that *Pfk* mRNA levels were comparable between *sea*^∆24^/*Df* mutants and *sea*^prec^/*Df* controls (Supplemental Figure S4).

Our gene expression studies suggest that CIC activity influences glycolytic enzyme activity at a post-transcriptional level. Considering that citrate is a key allosteric regulator of phosphofructokinase (PFK) (Pogson and Randle 1966; Tornheim and Lowenstein 1976; Kemp and Foe 1983; Usenik and Legiša 2010), and that *sea* mutant cells exhibit a significant depletion of cytosolic citrate (Morciano et al. 2009), decreased *Drosophila* CIC activity would be expected to induce elevated glycolytic flux. Consistent with this possibility, *sea*^∆24^/*Df* mutants that were fed a citrate-supplemented diet accumulated excess citrate and exhibited a significant decrease in pyruvate, lactate, and 2HG (Fig. 3A). Intriguingly, these results agree with the findings of a recent human case study, where a patient with a combined D-/L-2HGA exhibited decreased urinary 2HG levels and reduced cardiac symptoms following citrate treatment (Muhlhausen et al. 2014).

**Figure 3.**
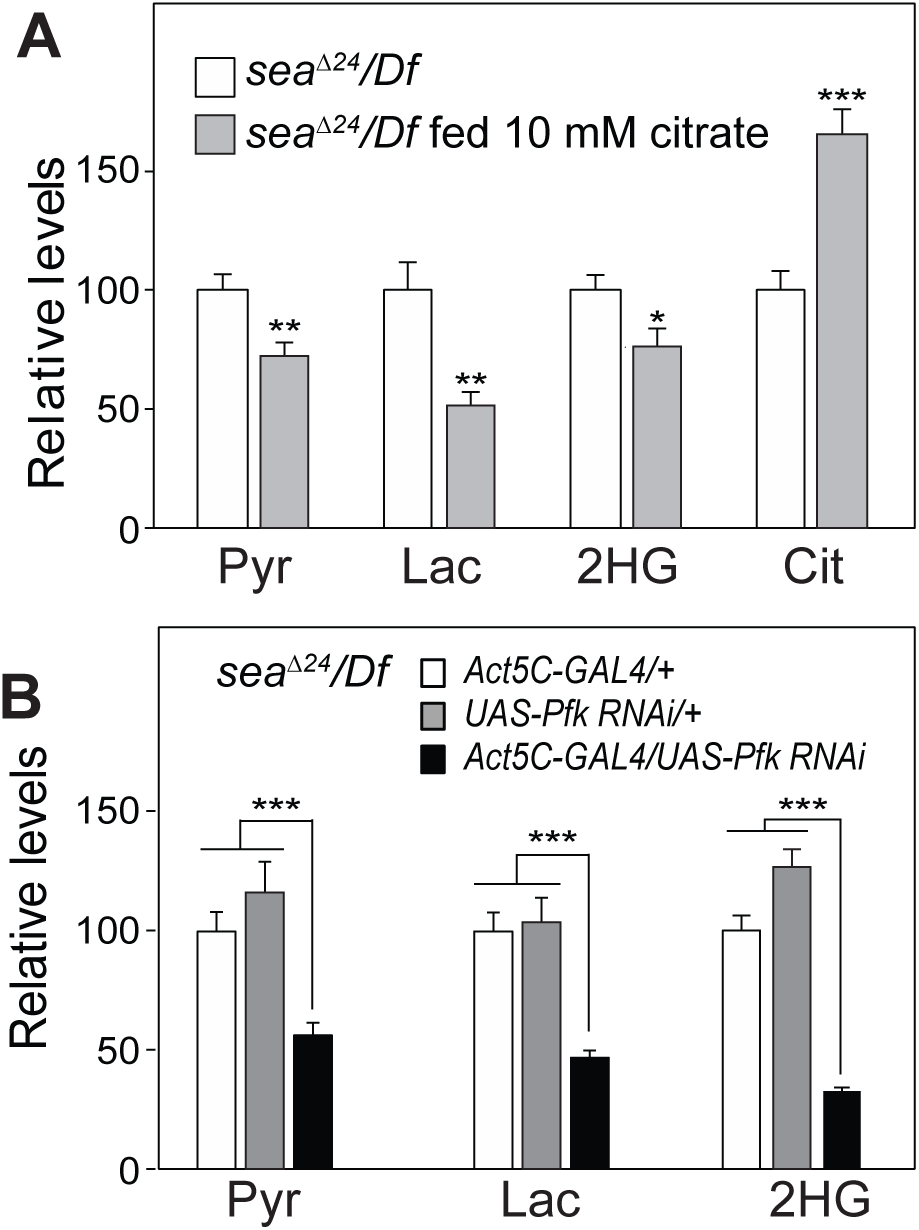
L-2HG levels in *sea* mutants are dependent on PFK activity. (A) *sea*^Δ24^/*Df* mutant larvae fed a semi-defined diet supplemented with 10 mM citrate accumulated excess citrate (cit) and displayed significant decreases in pyruvate (pyr), lactate (lac) and 2HG. (**B**) *Pfk-RNAi* reduces 2HG levels in *sea* mutant larvae. Data are shown as mean ± SEM, *n* = 6, \**P* < 0.05, \*\**P* < 0.01, \*\*\**P* < 0.001.

The ability of exogenous citrate to reduce steady-state levels of pyruvate, lactate, and L-2HG supports a model in which *sea*^∆24^/*Df* mutants accumulate excess L-2HG due to increased PFK activity. We further tested this hypothesis by using a *UAS-Pfk-RNAi* transgene to attenuate glycolysis in both control and *sea*^∆24^/*Df* mutant larvae. Ubiquitous expression of this transgene in a wild-type background reduced *Pfk* mRNA levels by 80% and induced a similar reduction in pyruvate, lactate, and 2HG levels, thereby confirming previous observations that, in larvae, these compounds are largely derived from glucose (Supplemental Fig. S5). Similarly, *Pfk-RNAi* expression in a *sea*^∆24^/*Df* mutant background induced an 80% decrease in *Pfk* mRNA levels, a 75% decrease in 2HG, and an ~50% decrease in pyruvate and lactate (Fig 3B and Supplemental Fig. S6). Overall, our experiments indicate that *sea* mutants accumulate excess L-2HG due to decreased cytosolic citrate levels, increased PFK activity, and elevated glycolytic flux.

#### sea mutants accumulate excess L-2HG due to decreased dL2HGDH activity

To understand how elevated glucose catabolism influences L-2HG metabolism, we used a genetic approach to determine if *sea* mutants display elevated L-2HG levels due to increased synthesis, decreased degradation, or a combination of both processes. We distinguished between these possibilities by measuring 2HG abundance in *dL2HGDH^12/14^*; *sea^∆24^*/*Df* double mutants, which are able to synthesize, but not degrade, L-2HG. If loss of CIC activity leads primarily to excess L-2HG synthesis, then *dL2HGDH^12/14^*; *sea^∆24^*/*Df* double mutants should accumulate more L-2HG than the *dL2HGDH*^12/14^ single mutant. In contrast, if *sea* mutants accumulate L-2HG due to decreased degradation, then L-2HG levels will be similar in both genetic backgrounds. GC-MS analysis revealed that double mutants harbored 2HG levels that were similar to those observed in single mutant controls (Fig. 4A), indicating that CIC activity primarily regulates L-2HG accumulation by controlling the degradation rate. In contrast, *dL2HGDH^12/14^; sea^∆24^*/*Df* double mutant larvae exhibited increased levels of lactate and pyruvate, as well as decreased citrate levels (Fig. 4A), suggesting that dL2HGDH does not influence these aspects of the *sea* mutant phenotype.

**Figure 4.**
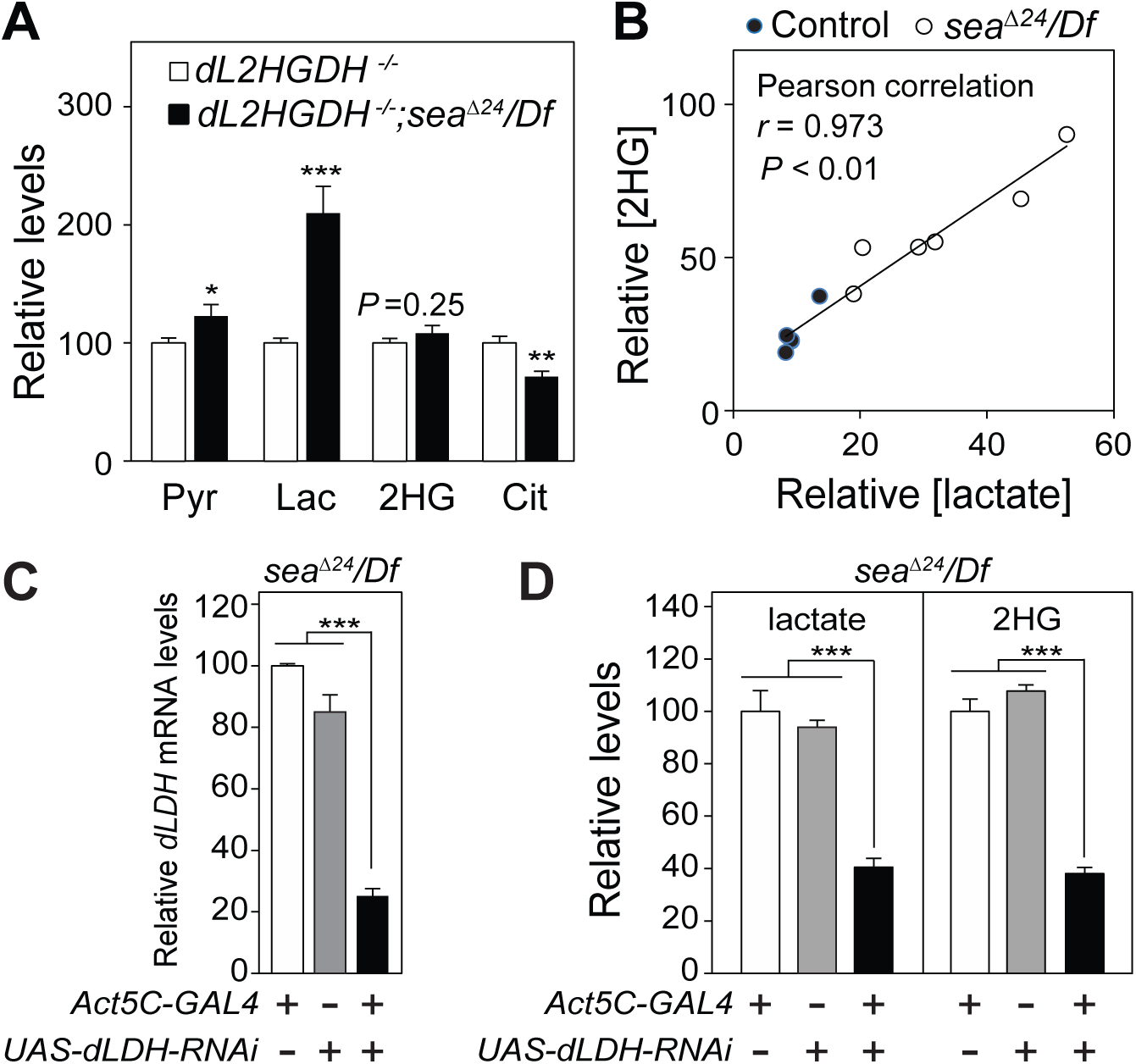
*sea* mutants accumulate excess L-2HG due to decreased degradation. (**A**) The relative abundance of pyruvate (pyr), lactate (lac), 2HG, and citrate (cit) in *dL2HGDH^12/14^* single mutants compared with *dL2HGDH^12/14^; sea*^∆24^/*Df* double mutants. Note that 2HG levels are similar in both strains. (**B**) Lactate and 2HG levels are highly correlated in individual larval samples. (**C**) Relative *dLdh* mRNA levels in *sea*^Δ24^/*Df* mutant larvae that ubiquitously express a *UAS-dLdh-RNAi* transgene. Data are shown as mean ± SEM, *n* = 3, \*\*\**P* < 0.001. (**D**) The relative abundance of lactate and 2HG in *sea*^Δ24^/*Df* mutant larvae that express the same *UAS-dLdh-RNAi* transgene used in panel (**C**). For (**A)** and **(D**), data are shown as mean ± SEM, *n* = 6, \**P* < 0.05, \*\**P* < 0.01, \*\*\**P* < 0.001.

Since we previously demonstrated that lactate production inhibits dL2HGDH activity and stabilizes the L-2HG pool (Li et al. 2017), our double mutant analysis hints at a model in which *sea^∆24^*/*Df* mutants accumulate excess L-2HG due to lactate-dependent dL2HGDH inhibition. Consistent with this possibility, qRT-PCR analysis revealed that *dL2HGDH* mRNA transcript levels are comparable between *sea^∆24^*/*Df* mutants and *sea^Prec^*/*Df* controls (Supplemental Fig. S7), indicating that CIC activity influences dL2HGDH activity at a post-transcriptional level. Moreover, we noticed a significant positive correlation between lactate and 2HG levels in *sea^∆24^*/*Df* mutants, *sea^Prec^*/*Df* controls, and *w^1118^*/*Df* controls (*r* = 0.973, *P* < 0.01; Fig. 4B, Supplemental Fig. S8). This observation suggests that lactate and L-2HG metabolism are coordinately regulated and supports a model in which elevated lactate levels stabilize the L-2HG pool. To directly test this hypothesis, we expressed a previously described *UAS*-*dLdh-RNAi* transgene in a *sea* mutant background. When compared with control strains, *dLdh-RNAi* induced an 80% decrease in *dLdh* mRNA levels and an ~60% reduction in both lactate and 2HG (Fig. 4C,D). Overall, these results support a model in which *sea* mutants primarily accumulate L-2HG due to a lactate-dependent decrease in dL2HGDH activity.

Finally, we examined the possibility that reduced CIC activity enhances the ability of dLDH to synthesize L-2HG, which could account for the increased rate of L-2HG synthesis observed in the ^13^C tracer experiments. qRT-PCR analysis revealed that *dLdh* gene expression is unchanged in *sea^∆24^*/*Df* mutants (Supplemental Figure S9), suggesting that any increase in dLDH activity must occur at an enzymatic level. Considering that acidic environments enhance the ability of mammalian LDHA to synthesize L-2HG (Nadtochiy et al. 2016; Teng et al. 2016; Intlekofer et al. 2017), the excess lactate present within *sea^∆24^*/*Df* mutants might promote L-2HG synthesis by lowering the pH of larval tissues. In support of this hypothesis, acidic pH enhanced the ability of purified dLDH to catalyze the formation of L-2HG from 2-OG *in vitro* (Fig. 5, Supplemental Fig. S10), indicating that increased lactate production could promote L-2HG synthesis. Therefore, while decreased degradation appears to be the primary mechanism responsible for expansion of the L-2HG pool in *sea* mutants, changes in intracellular pH might also contribute to this phenotype.

**Figure 5.**
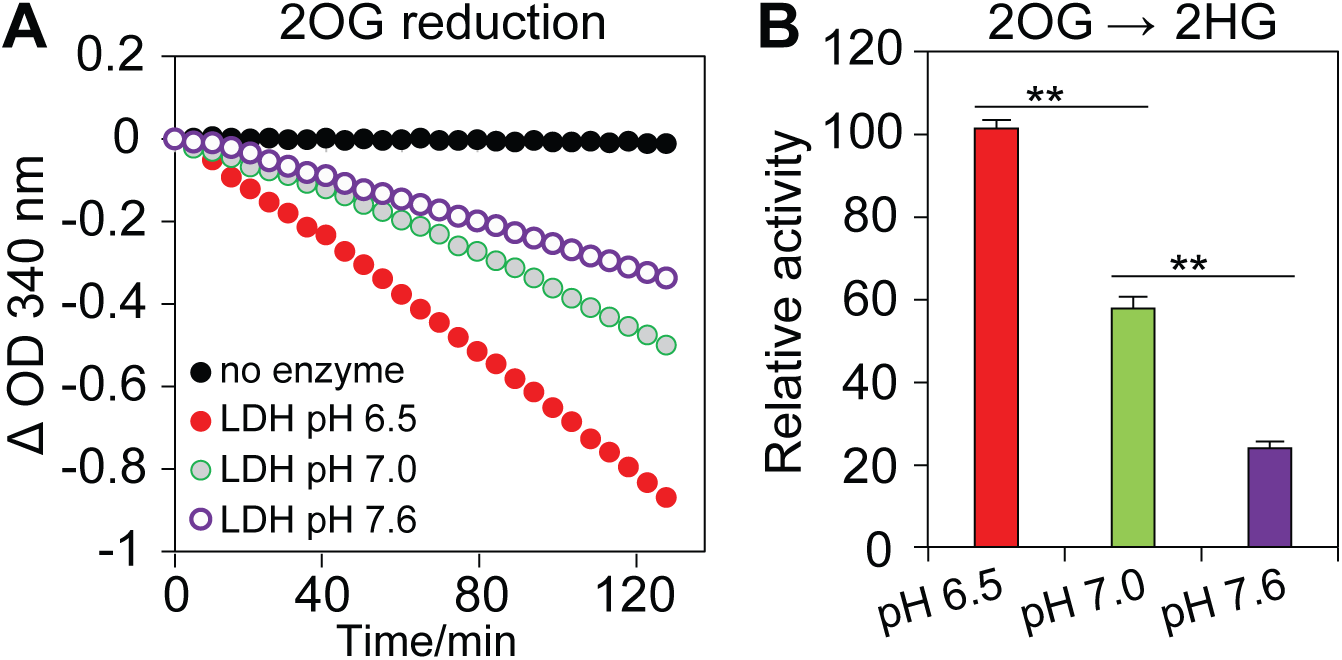
The effects of pH on the dLDH catalyzed formation of L-2HG from 2OG. (**A**) Purified *Drosophila* LDH was incubated with 2OG, NADH, pH adjusted buffers. The reaction rates were measured by changes in NADH concentration (absorbance at 340 nm). (**B**) Relative dLDH activity at different pH values was calculated based on the slopes in panel A. Data are shown as mean ± SD, *n* = 3, \*\**P* < 0.01.

#### L-2HG accumulation is coordinately regulated by glycolysis and the TCA cycle

The CIC plays a central role in cellular metabolism by controlling the amount of citrate that exits the TCA cycle and enters the cytosol. This function serves many purposes in cellular physiology, such as providing substrate for fatty acid synthesis, controlling histone acetylation, and regulating cellular redox balance (Palmieri 2004; Morciano et al. 2009; Palmieri 2013; Dolce et al. 2014). In addition, cytosolic citrate is an allosteric regulator of PFK, which represents one of three rate-limiting glycolytic enzymes (Pogson and Randle 1966; Tornheim and Lowenstein 1976; Kemp and Foe 1983; Usenik and Legiša 2010). This feedback mechanism fine-tunes central carbon metabolism by serving as a signal to slow glycolysis during times of sufficient energy production. Our findings suggest that the role of citrate as an allosteric regulator of PFK represents the primary mechanism that induces L-2HG accumulation in *sea* mutants.

Based on both our study and previous observations in human cell culture (Jiang et al. 2017), CIC deficiency induces elevated glycolytic flux and decreased citrate production (see Supplemental Figure S11), thereby generating a cellular environment in which pyruvate levels rise due to both increased production and decreased TCA cycle utilization. As observed in *Drosophila*, human patients, and CIC-deficient cells (Nota et al. 2013; Prasun et al. 2015; Jiang et al. 2017; Li and Tennessen 2017), these metabolic disruptions result in enhanced lactate synthesis, which, at least in the fly, inhibits dL2HGDH and stabilizes the L-2HG pool. Such a model would also explain why citrate treatment could reduce 2HG levels in a patient with combined D-/L-2HGA (Muhlhausen et al. 2014), as partial restoration of cytosolic citrate levels would inhibit PFK and reduce lactate production. In addition, our studies in *Drosophila* suggest that an LDH inhibitor, such as oxamate, could also alleviate some of the symptoms associated with combined D-/L-2HGA.

Our findings also highlight the role of dL2HGDH in controlling L-2HG accumulation. While dLDH synthesizes most of the larval L-2HG pool, the kinetics of this reaction are poor and the only reasonable explanation for how flies accumulate such high L-2HG levels rests upon our observations that dL2HGDH activity is sensitive to lactate (Li et al. 2017). Since larval metabolism is highly glycolytic, aerobic lactate production stabilizes the L-2HG pool and allows for dramatic accumulation of this metabolite. Whether this mechanism also functions in human cells warrants further examination, although the positive correlation between L-2HG and lactate has been repeatedly observed in mammals. Notably, elevated L-2HG levels are associated with increased lactate production in mouse CD8^+^ T cells, human cells with disrupted 2OG metabolism, and in mammalian cells subjected to hypoxic conditions (Intlekofer et al. 2015; Oldham et al. 2015; Burr et al. 2016; Tyrakis et al. 2016). Furthermore, we would also highlight a somewhat overlooked observation that mouse L2HGDH activity is inhibited by acidic pH (Nadtochiy et al. 2016), suggesting that even if lactate doesn’t directly regulate L-2HG degradation, excess lactate accumulation could establish a microenvironment that stabilizes the L-2HG pool. When considered in this context, our findings hint at a conserved feed-forward loop in animal metabolism that links lactate synthesis with L-2HG accumulation.

Finally, our study raises the question of why D-2HG levels remain low in *sea* mutants. After all, D-2HG levels exceed those of L-2HG in both combined D-/L-2HGA patients and CIC-deficient cells. While an adequate explanation requires a more detailed examination of *Drosophila* D-2HG metabolism, this discrepancy highlights a key difference between fly and human metabolism. Throughout our analyses, we repeatedly observed that flies accumulate minimal amounts of D-2HG, suggesting that the metabolic enzymes driving D-2HG accumulation in humans have either diverged in flies such that they no longer synthesize this molecule or that D-2HG is only produced under specific cellular conditions. When considered in light of these findings, our findings highlight the importance of L-2HG in cellular metabolism, as the mechanisms that control L-2HG accumulation are conserved across phyla. Moreover, these observations reinforce the notion that *Drosophila* genetics provides a powerful tool for dissecting the metabolic mechanisms that underlie L-2HG metabolism.

## Materials and Methods

### Drosophila husbandry and genetics

Fly stocks were maintained on standard Bloomington stock center media. Larvae were raised on molasses agar plates with yeast paste spread on the surface. Both the *sea* mutants (*sea^∆24^*) and the precise excision control strain (*sea^Prec^*; previously noted as *Rev^24^*) were kindly provided by Dr. Giovanni Cenci (Morciano et al. 2009). All the experiments used a trans-heterozygous combination of *sea^∆24^* and a molecularly-defined deficiency, ando all controls consisted of a trans-heterozygous combination of *sea*^*Prec*^ and the same molecularly-defined deficiency. A complete list of BDSC stocks used in this study are available in Supplemental Table S3. The *UAS*-*sea* strain was generated by injecting the DGRC plasmid UFO06122 into BDSC Stock 8621. *dL2HGDH* mutant strains are the same as reported previously (Li et al. 2017).

### Metabolomics and metabolic flux analysis

Middle third instar (mid-L3) larval samples were prepared and analyzed using GC-MS as described previously (Li et al. 2017). Spectral data preprocessing was performed using MetAlign software (Lommen 2009). For metabolic flux measurements, mid-L3 larvae were fed with Semi-defined Medium (Backhaus et al. 1984) containing 50% D-glucose-^13^C_6_ for 2 hours, then metabolites were detected using GC-MS. The isotopologue distributions were corrected based on the natural abundance of elements. The metabolic flux *f*_*x*_ was estimated based on the formula *X*^*L*^/*X*^*T*^ = *p*(1–*exp*(-*f*_*x*_\**t*/*X*^*T*^)), where *X*^*L*^ is the amount of ^13^C labeled metabolite, *X*^*T*^ is the amount of total metabolite pool, *p* is the percentage of glucose-^13^C_6_.

### qRT–PCR

Total RNAs were extracted using Trizol reagent (ThermoFisher Scientific). cDNA was made using Maxima First Strand cDNA Synthesis Kit (ThermoFisher Scientific), and qPCR was performed using FastStart Essential DNA Green Master Kit (Roche Diagnostics) in a LightCycler 96 instrument (Roche Diagnostics). The primers for *rp49* and *LDH* are the same as reported previously (Li et al. 2017). Additional primer sequences are described in Supplemental Table S4. mRNA levels were normalized to *rp49*.

### Enzyme activity assay

*Drosophila* LDH was purified as described previously (Li et al. 2017). The assay was performed in 100 mM PBS at 25 °C with the indicated pH value. Each reaction contains 10 mM 2OG and 1 mM NADH (Sigma-Aldrich). The values of OD_340 nm_ were recorded by a Cytation 3 plate reader (BioTek).

### Statistical analysis

Multivariate data analysis (principal component analysis, PCA) was performed using MetaboAnalyst (Xia and Wishart 2016). Unless noted, two-tailed Student’s *t*-test was performed to do univariate statistical analysis and Bonferroni correction was used for multiple comparisons.

### RNA-seq analysis

RNA was purified from staged mid-L3 larvae using a RNeasy Mini Kit (Qiagen). Sequencing was performed using an Illumina NextSeq500 platform with 75 bp sequencing module generating 41 bp paired-end reads. After the sequencing run, demultiplexing with performed with bcl2fastq v2.20.0.422. All files were trimmed using cutadapt v1.12 with the parameters: “-a AGATCGGAAGAGC -m 30 -q 30” (Martin 2011). Remaining read pairs were mapped against Flybase 6.19 using hisat2 v2.1.0 with “--no-unal --no-mixed” parameters (Kim et al. 2015). Gene counts were produced using HTseq-Count v0.9.0 (Anders et al. 2015). The R package DESeq2 was used to identify the differentially expressed genes (Love et al. 2014).

## Acknowledgments

We thank the Bloomington *Drosophila* Stock Center, the *Drosophila* Genomics Resource Center, Flybase, and Center for Genomics and Bioinformatics at Indiana University. We also thank Dr. Giovanni Cenci (University of Rome) for strains and helpful advice. This project was supported by an Indiana CTSI award, which was funded in part by NIH grant UL1TR001108. J.M.T. is supported by NIH R35 Maximizing Investigators’ Research Award 1R35GM119557.

## Author Contributions

H.L. and J.M.T. designed this study, wrote the manuscript, and generated figures. H.L. and A.J.H. generated reagents and conducted the experiments.

---

**Supplemental Figure S1.**
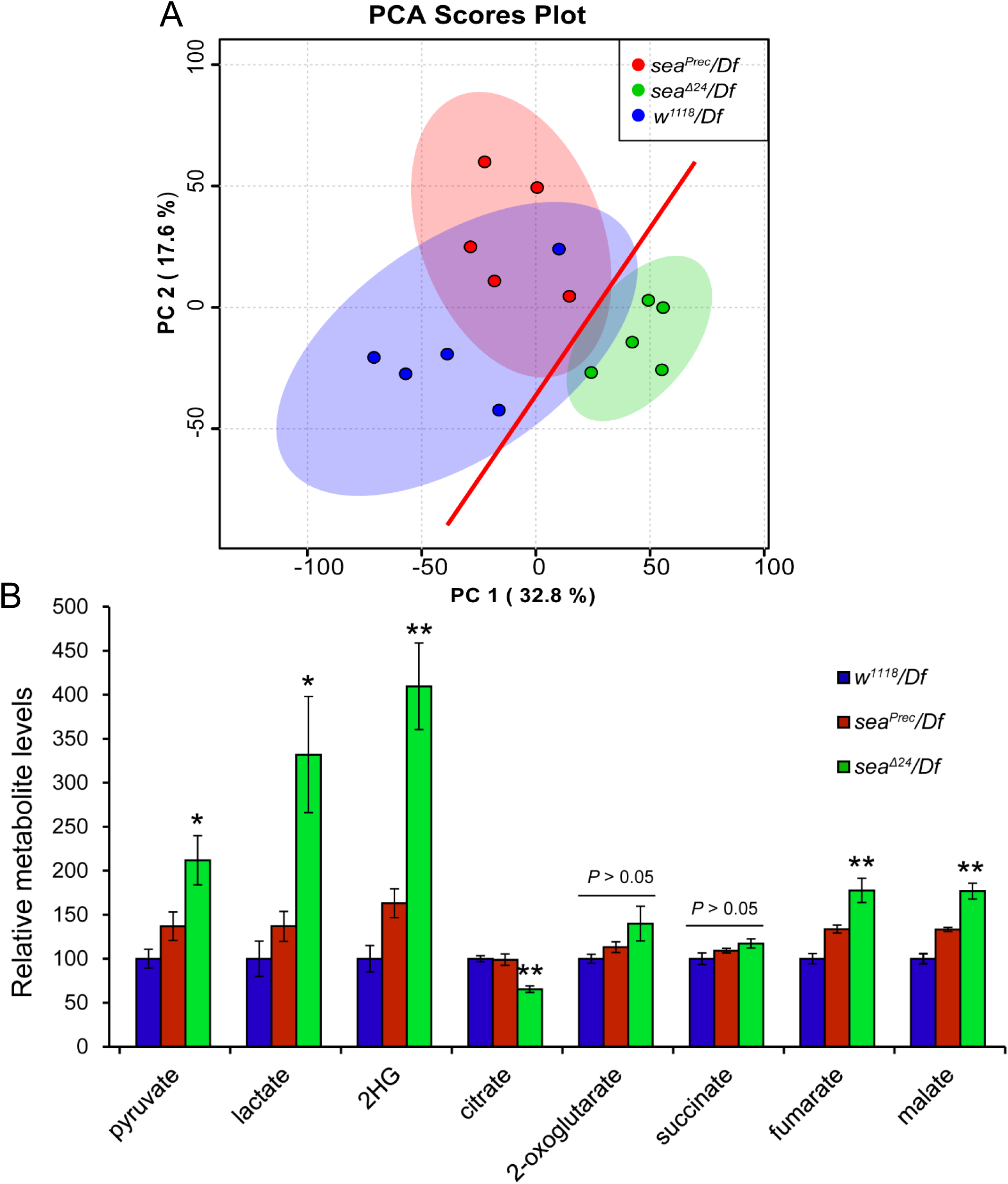
The metabolic changes caused by sea deficiency in *Drosophila* larvae. (**A**) PCA scores plots of metabolic profiles of sea mutants (*sea*^Δ24^/*Df*) and two control strains (*sea*^Prec^/*Df* and *w*^1118^/*Df*). (**B**) A comparison of relative metabolite levels between sea mutant larvae and the controls. *Df* refers to the molecularly-define deficiency *Df(3R)Exel8153*, which uncovers the sea gene. Data shown as mean ± SEM, *n* = 5, \**P* < 0.05, \*\**P* < 0.01.

**Supplemental Figure S2.**
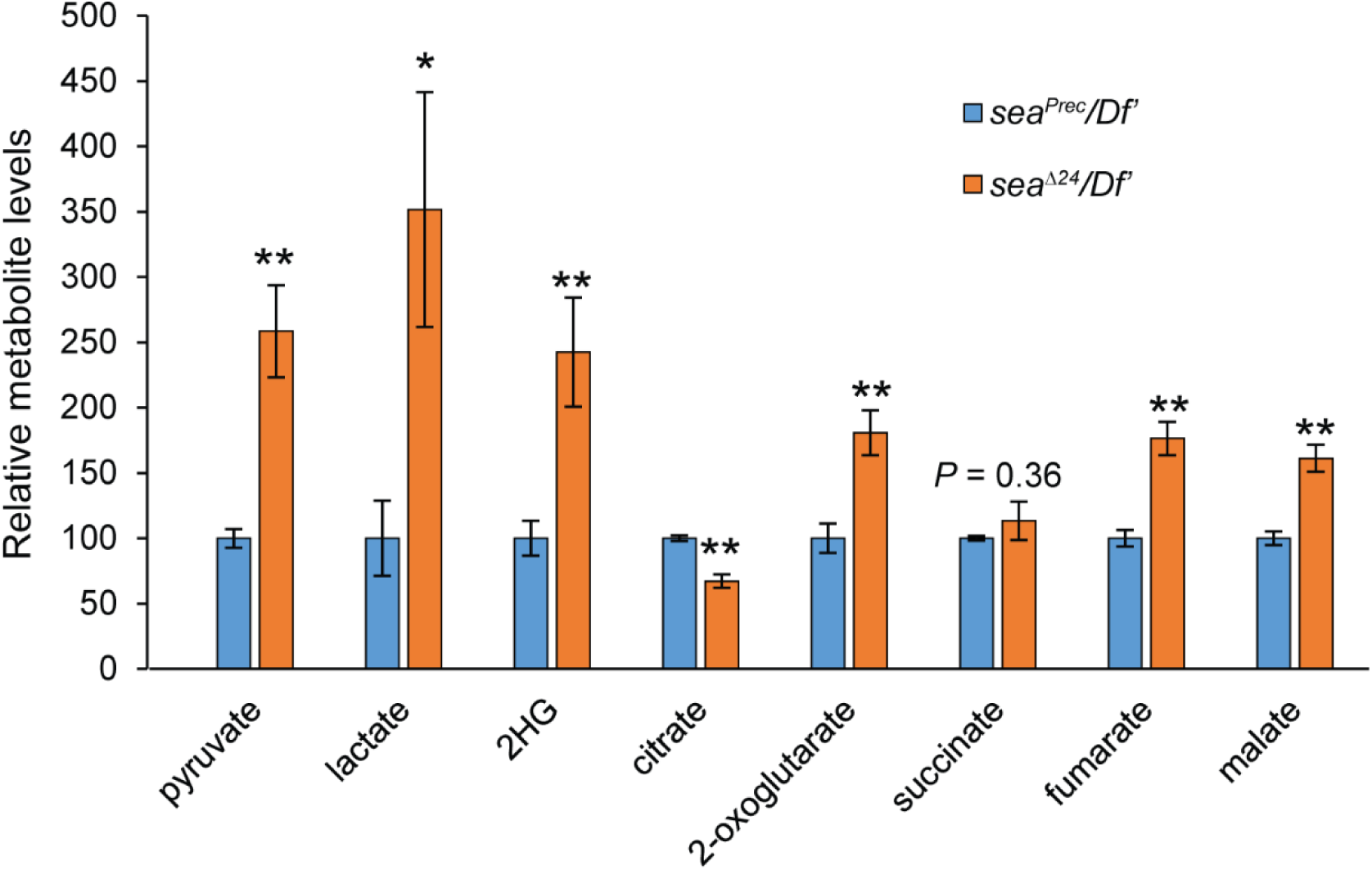
Metabolic defects induced by the *sea*^Δ24^ mutation were confirmed using a different deficiency strain *Df(3R)BSC469* (*Df’*). Data are shown as mean ± SEM. *n* = 6, \**P* < 0.05, \*\**P* < 0.01.

**Supplemental Figure S3.**
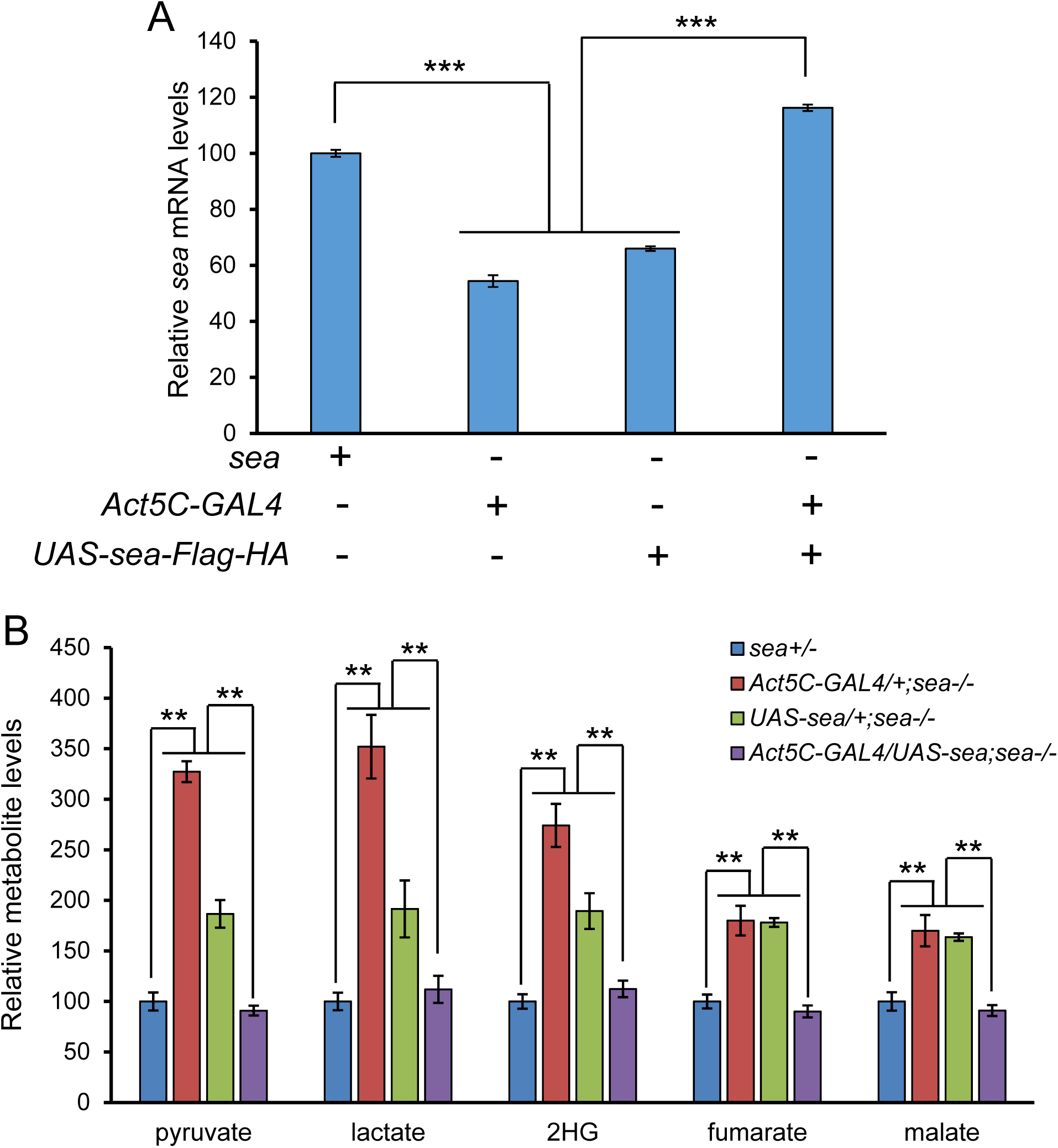
*GAL4*-driven expression of *UAS*-*sea-Flag-HA* rescued the phenotypes caused by the *sea* mutation. (**A**) Relative *sea* mRNA levels in indicated genotypes. Data shown as mean ± SEM, *n* = 4, \*\*\**P* < 0.001. (**B**) Relative metabolite levels in indicated genotypes. Data shown as mean ± SEM, *n* = 6, \*\**P* < 0.01.

**Supplemental Figure S4.**
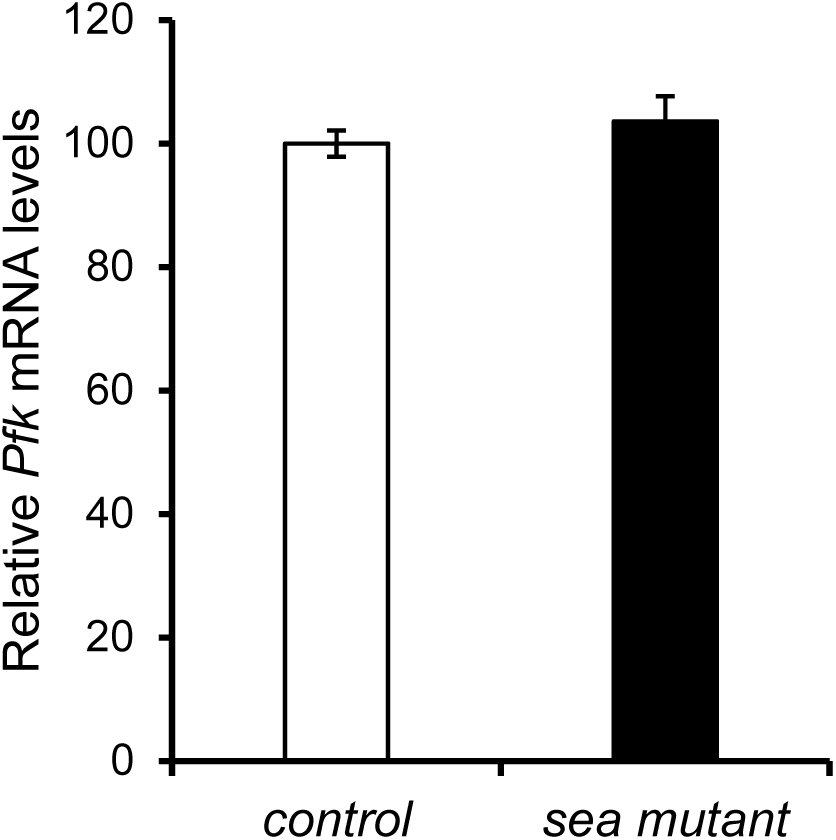
A comparison of *Pfk* mRNA levels between the *sea* mutant (*sea*^Δ24^/*Df*) and control (*sea*^Prec^/*Df*). Data shown as mean ± SEM, *n* = 3, *P* > 0.05.

**Supplemental Figure S5.**
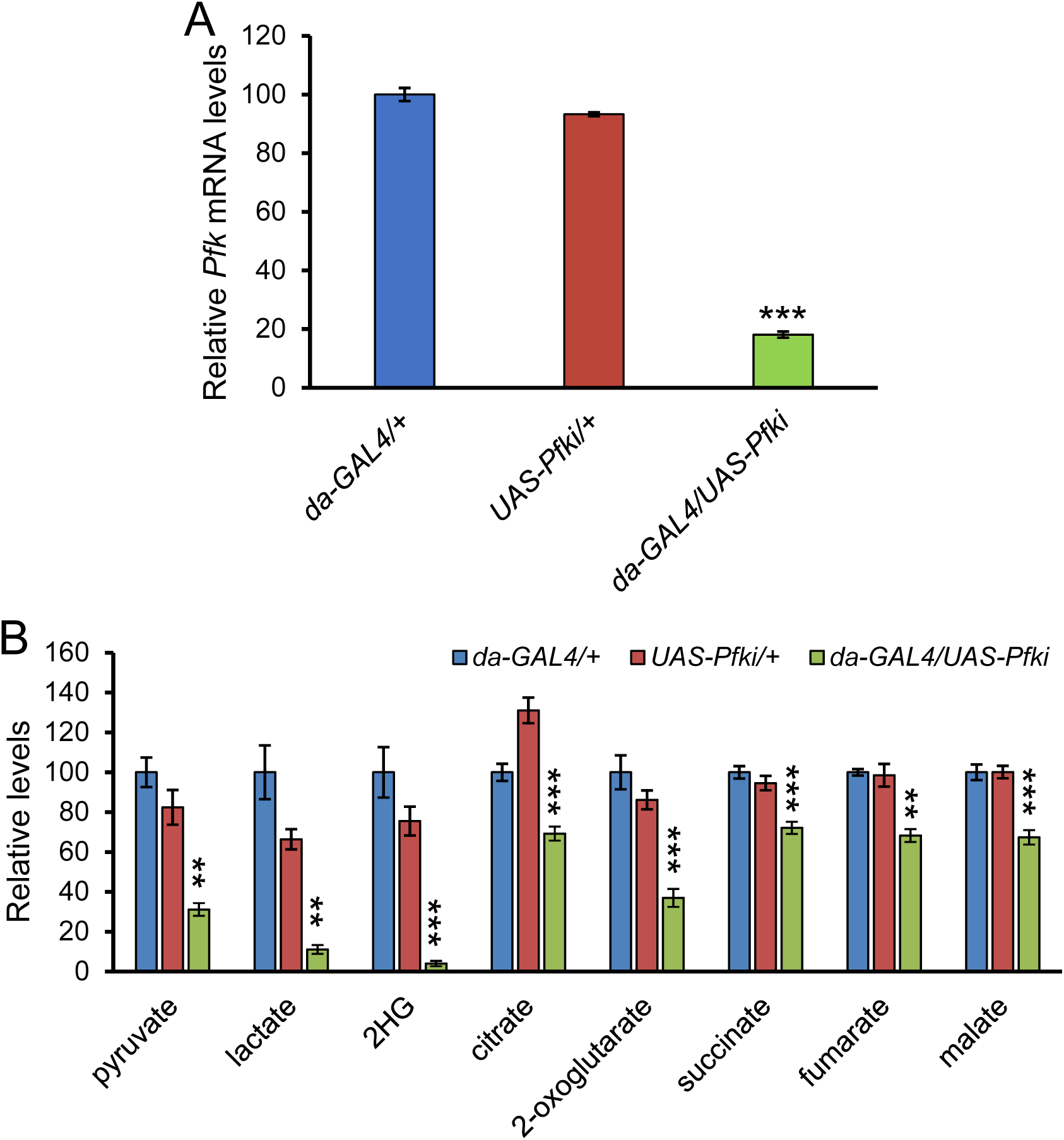
Changed metabolic profiles caused by *Pfk* knockdown. (**A**) Expression of dsRNA for RNAi of *Pfk* (*Pfki*) down-regulates the transcriptional levels of *Pfk* significantly. Data shown as mean ± SEM, *n* = 3, \*\*\**P* < 0.001. (**B**) Changed metabolites induced by *Pfk* knockdown. Data shown as mean ± SEM, *n* = 6, \*\**P* < 0.01, \*\*\**P* < 0.001.

**Supplemental Figure S6.**
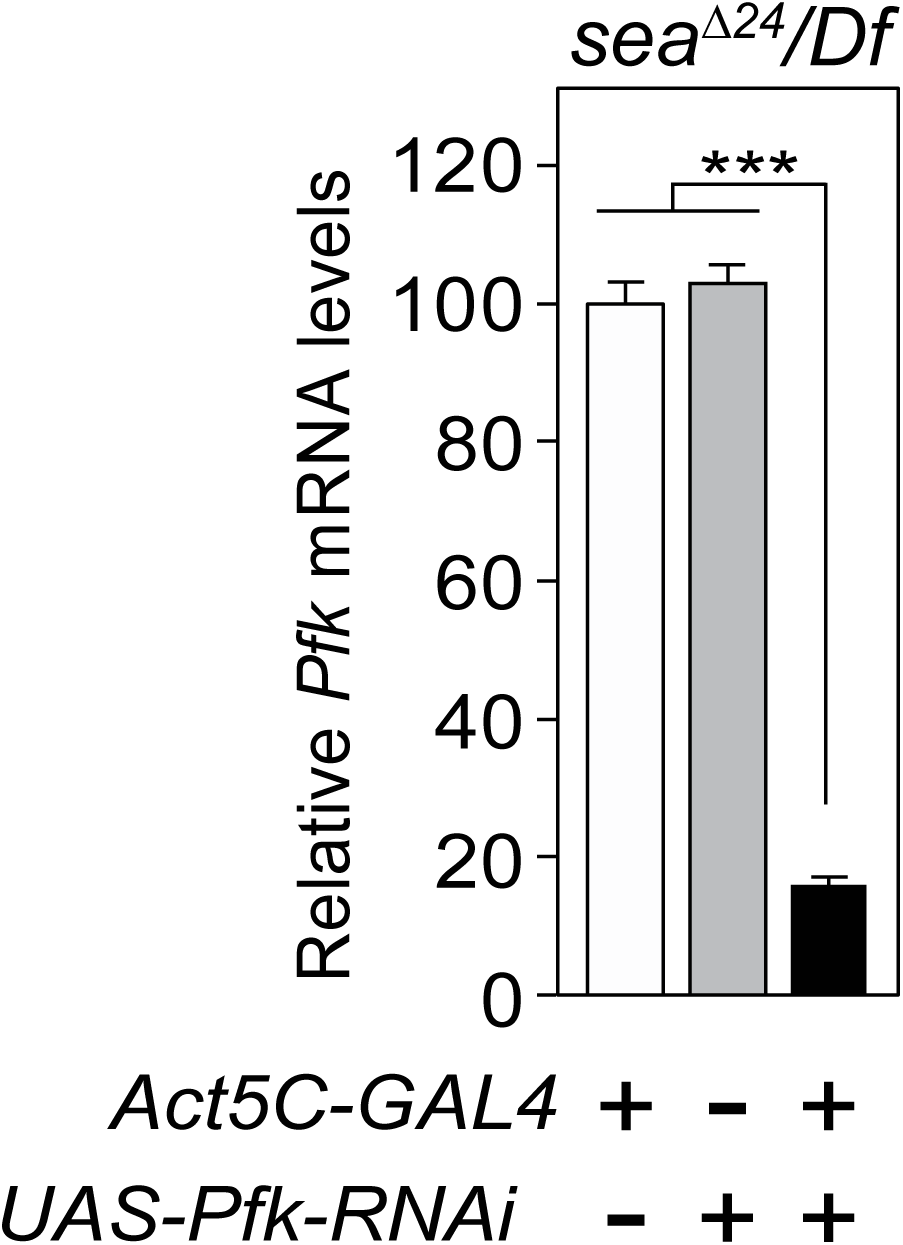
The relative mRNA levels of *Pfk* in *sea* mutant larvae with indicated genotypes. Data are shown as mean ± SEM, *n* = 3, \*\*\**P* < 0.001.

**Supplemental Figure S7.**
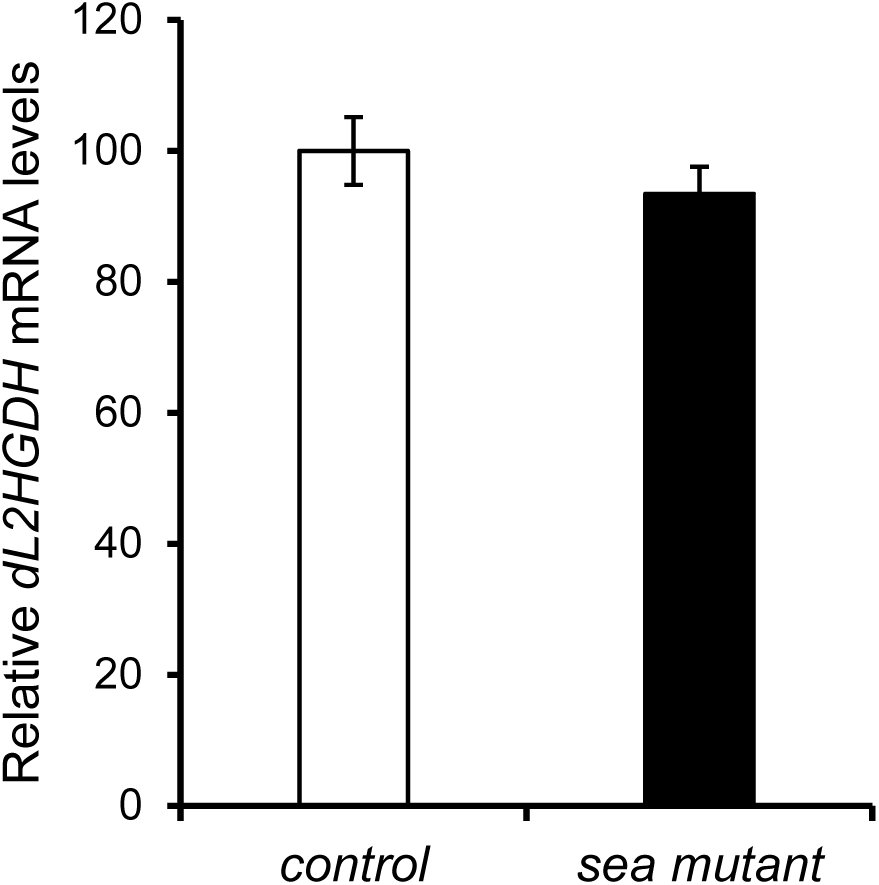
A comparison of *dL2HGDH* mRNA levels between the *sea* mutant (*sea*^Δ24^/*Df*) and control (*sea*^Prec^/*Df*). Data shown as mean ± SEM, *n* = 3, *P* > 0.05.

**Supplemental Figure S8.**
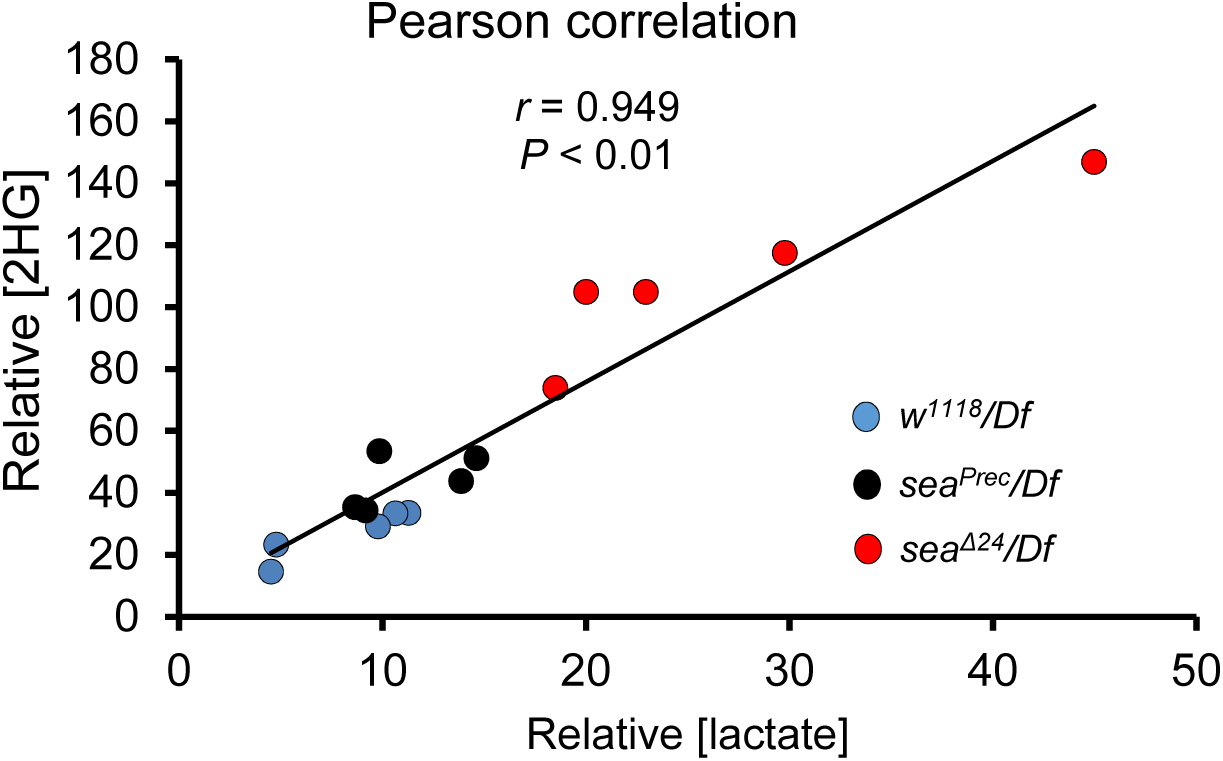
The correlation between the levels of lactate and 2HG in individual mL3 larval sample from different groups. *sea* mutant (*sea*^Δ24^/*Df*) and its two controls (*sea*^Prec^/*Df* and *w^1118^/Df*). *Df* refers to the molecularly-define deficiency *Df(3R)Exel8153*, which uncovers the *sea* gene.

**Supplemental Figure S9.**
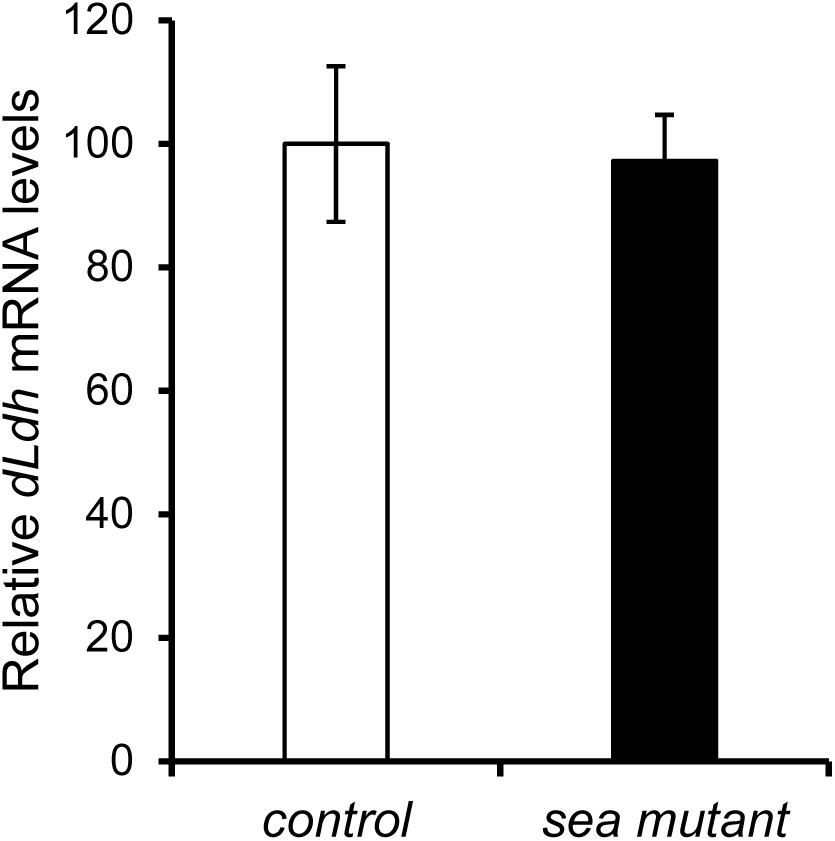
A comparison of *dLdh* mRNA levels between the *sea* mutant (*sea*^Δ24^/*Df*) and control (*sea*^Prec^/*Df*). Data shown as mean ± SEM, *n* = 3, *P* > 0.05.

**Supplemental Figure S10.**
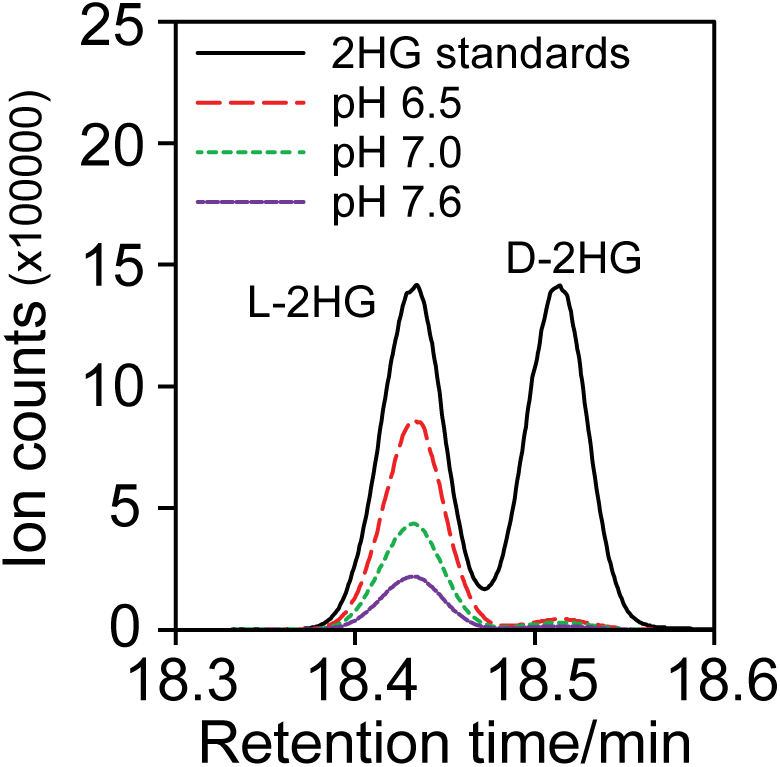
The products of each reaction were detected by GC-MS with chiral derivatization to confirm that the dominant product is L-2HG.

**Supplemental Figure S11.**
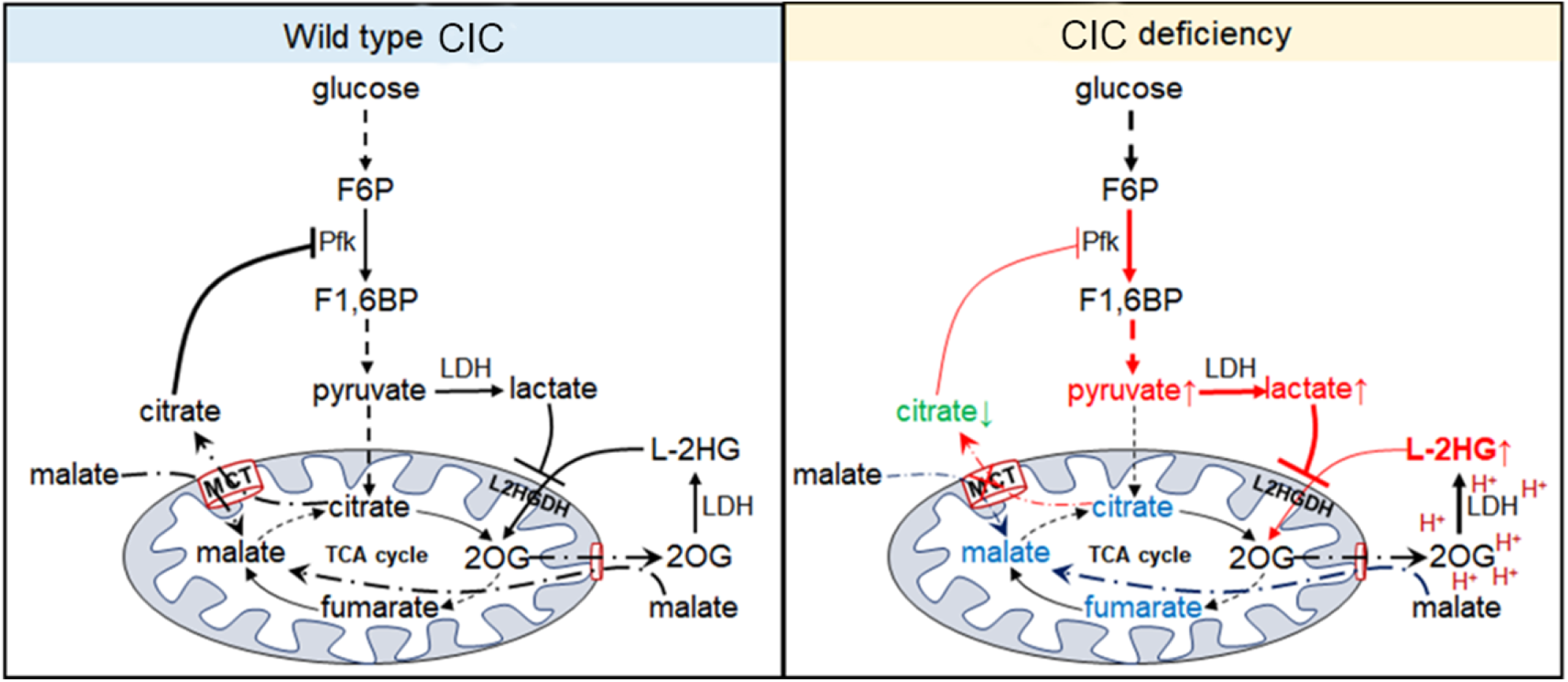
Schematic summary illustrating the mechanism by which mitochondrial citrate transporter (CIC) affects the accumulation of L-2HG.

**Supplemental Table S2.** The expression of genes encoding glycolytic enzymes were compared between *sea[24]/Df mutants and sea[prec]/Df* controls. Data derived from Supplemental Table S1.

| Enzyme Function | FBgn | Gene | Base Mean |
| --- | --- | --- | --- |
| hexokinase (best match to human hexokinase IV) | FBgn0001186 | Hex-A | 16053.9055 |
| hexokinase | FBgn0001187 | Hex-C | 3978.6868 |
| hexokinase | FBgn0042710 | Hex-t2 | 135.551903 |
| hexokinase | FBgn0042711 | Hex-t1 | 18.6612123 |
| phosphoglucose isomerase | FBgn0003074 | Pgi | 68381.3879 |
| Phosphofructokinase | FBgn0003071 | Pfk | 36224.9971 |
| aldolase | FBgn0000064 | Ald | 264899.634 |
| aldolase | FBgn0039425 | CG5432 | 142.641361 |
| triose phosphate isomerase | FBgn0086355 | Tpi | 52056.7575 |
| glycerol 3 phosphate dehydrogenase | FBgn0001128 | Gpdh | 75417.8956 |
| glyceraldehyde phosphate dehydrogenase | FBgn0001091 | Gapdh1 | 161678.953 |
| glyceraldehyde phosphate dehydrogenase | FBgn0001092 | Gapdh2 | 257663.529 |
| glyceraldehyde 3-phosphate dehydrogenase | FBgn0034173 | CG9010 | 89.5456135 |
| phosphoglycerate kinase | FBgn0250906 | Pgk | 45043.1381 |
| phosphoglycerate kinase | FBgn0031451 | CG9961 | 149.983451 |
| phosphoglycerate mutase | FBgn0014869 | Pglym78 | 55187.6462 |
| phosphoglycerate mutase | FBgn0011270 | Pglym87 | 65.5693506 |
| phosphoglycerate mutase | FBgn0038957 | CG7059 | 190.487025 |
| enolase | FBgn0000579 | Eno | 183715.725 |
| pyruvate kinase | FBgn0267385 | PyK | 77869.845 |
| pyruvate kinase | FBgn0038258 | CG7362 | 16.1795929 |
| pyruvate kinase | FBgn0038952 | CG7069 | 265.137004 |
| pyruvate kinase | FBgn0031462 | CG2964 | 64.1806881 |
| pyruvate kinase | FBgn0036723 | CG12229 | 123.000086 |
| L-lactate dehydrogenase | FBgn0001258 | dLdh (ImpL3) | 36449.7937 |
| L-lactate dehydrogenase | FBgn0033856 | CG13334 | 11.7465735 |

*a[24]/Df* mutants and *sea[prec]/Df* controls. Data derived from Supplemental Table S1.
| log2 Fold Change | Adj. pvalue | C1 | C2 | C3 | S1 | S2 |
| --- | --- | --- | --- | --- | --- | --- |
| 0.253696585 | 0.00778946 | 15012.994 | 15092.9436 | 13832.9756 | 17121.6912 | 19123.5513 |
| 0.872410777 | 3.2301E-17 | 2889.93847 | 2725.51689 | 2817.52654 | 5470.20739 | 4504.23521 |
| -1.729302381 | 0.00576783 | 219.285272 | 203.498449 | 202.707079 | 91.7820033 | 81.6830219 |
| -1.474911387 | 0.12826018 | 33.1945596 | 17.8806169 | 31.5774847 | 12.5596426 | 15.5586708 |
| -0.03221151 | 0.76081911 | 67100.2874 | 70343.1982 | 69987.9297 | 66521.6637 | 66129.2131 |
| -0.052785462 | 0.62601693 | 36943.533 | 35739.0958 | 37978.5464 | 32994.181 | 35264.6999 |
| -0.256211007 | 0.01078449 | 288390.31 | 270970.531 | 305717.928 | 222156.89 | 237776.359 |
| -1.568536847 | 0.01547101 | 265.556477 | 159.222636 | 215.94925 | 103.375519 | 91.4071912 |
| 0.518097535 | 0.00105756 | 43143.8744 | 44734.749 | 40545.4904 | 51883.8834 | 56379.761 |
| 0.092188272 | 0.40844882 | 71573.5058 | 73249.2242 | 74205.0518 | 84959.219 | 71945.2388 |
| -0.092618061 | 0.42659743 | 165872.208 | 165335.253 | 169390.796 | 143081.38 | 149804.717 |
| -0.004561474 | 0.82612933 | 259528.143 | 250984.259 | 263698.482 | 238121.162 | 256667.503 |
| -1.675314663 | 0.00385912 | 136.801821 | 114.095365 | 158.906052 | 66.6627182 | 47.6484294 |
| 0.1838583 | 0.15604251 | 41779.8798 | 42322.5687 | 42425.8786 | 42844.8052 | 45229.0561 |
| -1.199485062 | 0.00047888 | 219.285272 | 166.0343 | 242.433592 | 118.833541 | 87.5175235 |
| -0.03245939 | 0.70724869 | 55586.8047 | 55005.0348 | 56831.3234 | 49367.1242 | 50487.8869 |
| -2.389610059 | 7.3359E-05 | 122.719281 | 80.8885049 | 127.328567 | 19.322527 | 35.9794263 |
| -0.348319072 | 0.30505717 | 227.332438 | 229.042188 | 184.371766 | 193.22527 | 185.731633 |
| 0.073416727 | 0.62075545 | 171966.931 | 186152.548 | 179004.612 | 169255.675 | 188655.691 |
| -0.067375215 | 0.44664918 | 78039.4037 | 81375.5389 | 79645.5468 | 70544.6138 | 73657.665 |
| -2.948149931 | 0.00041598 | 25.1473936 | 26.3951963 | 34.6333703 | 2.89837905 | 6.80691849 |
| -0.815450539 | 0.01028556 | 407.387777 | 264.803421 | 343.277818 | 223.175187 | 211.014473 |
| -1.487889218 | 0.05352542 | 93.5483043 | 91.9574582 | 98.8069683 | 43.4756858 | 52.5105141 |
| -2.142221376 | 0.00014932 | 215.261689 | 176.251795 | 210.856108 | 42.5095594 | 78.7657711 |
| -0.256217906 | 0.06355449 | 40336.4194 | 39729.0278 | 38966.6161 | 27624.4507 | 33914.9852 |
| -2.036471702 | 0.02561563 | 20.1179149 | 17.0291589 | 19.3539422 | 0.96612635 | 5.83450156 |

| S3 | C Mean | S Mean |
| --- | --- | --- |
| 16139.2772 | 14646.3044 | 17461.5065 |
| 5464.69632 | 2810.99397 | 5146.37964 |
| 14.3555945 | 208.496934 | 62.6068732 |
| 1.19629954 | 27.5508871 | 9.77153764 |
| 70206.0351 | 69143.8051 | 67618.9706 |
| 38429.9266 | 36887.0584 | 35562.9358 |
| 264385.788 | 288359.59 | 241439.679 |
| 20.3370923 | 213.576121 | 71.706601 |
| 75652.7869 | 42808.0379 | 61305.4771 |
| 76575.1339 | 73009.2606 | 77826.5305 |
| 176589.365 | 166866.085 | 156491.821 |
| 276981.626 | 258070.295 | 257256.764 |
| 13.159295 | 136.601079 | 42.4901475 |
| 55656.64 | 42176.109 | 47910.1671 |
| 65.796475 | 209.251055 | 90.7158465 |
| 63847.703 | 55807.721 | 54567.5714 |
| 7.17779727 | 110.312118 | 20.8265835 |
| 123.218853 | 213.58213 | 167.391919 |
| 207258.896 | 179041.363 | 188390.087 |
| 83956.3021 | 79686.8298 | 76052.8603 |
| 1.19629954 | 28.7253201 | 3.6338657 |
| 141.163346 | 338.489672 | 191.784335 |
| 4.78519818 | 94.7709102 | 33.590466 |
| 14.3555945 | 200.789864 | 45.2103084 |
| 38127.2628 | 39677.3544 | 33222.2329 |
| 7.17779727 | 18.833672 | 4.65947506 |

**Supplemental Table 3.**
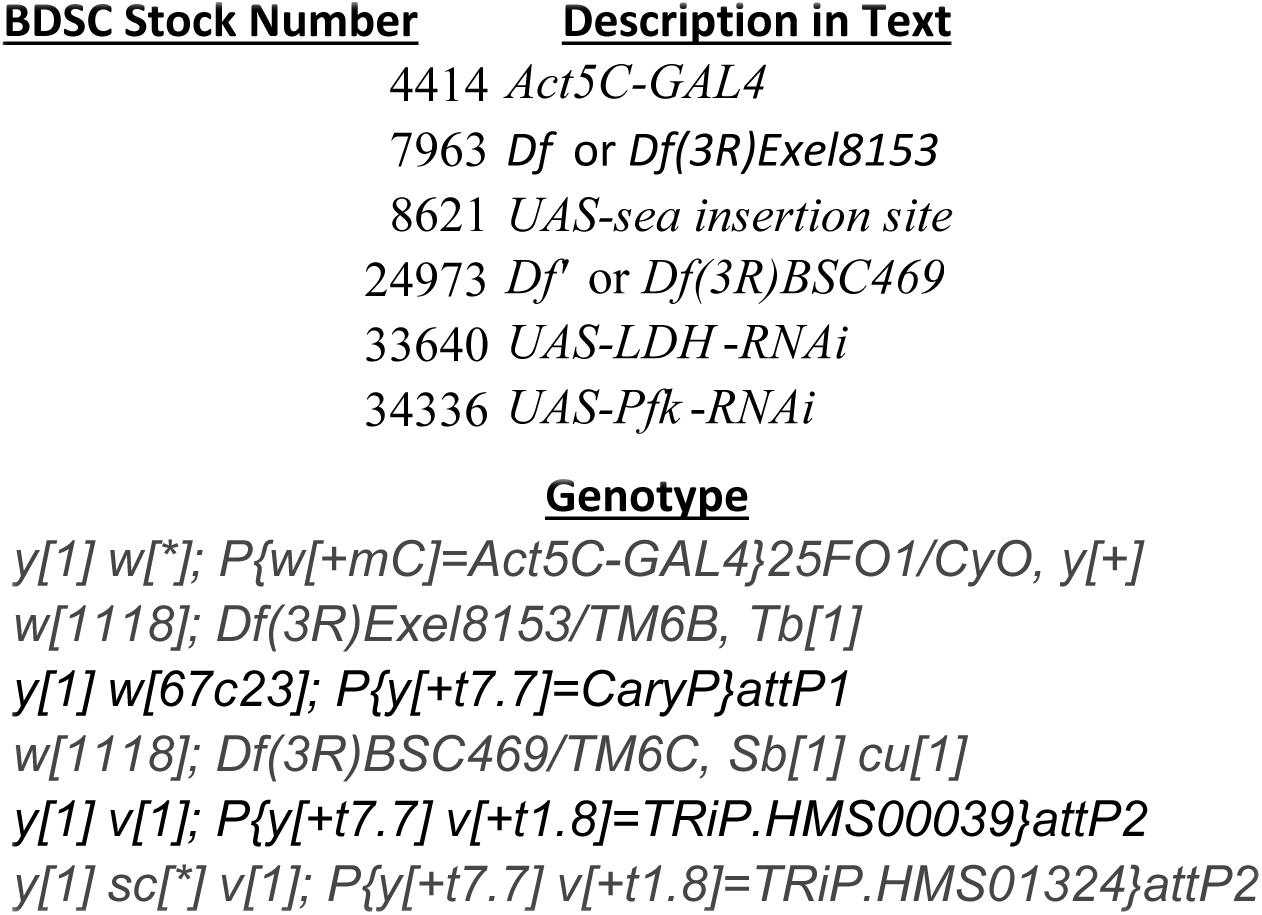
BDSC strains used for genetic analysis

**Supplemental Table 4.**
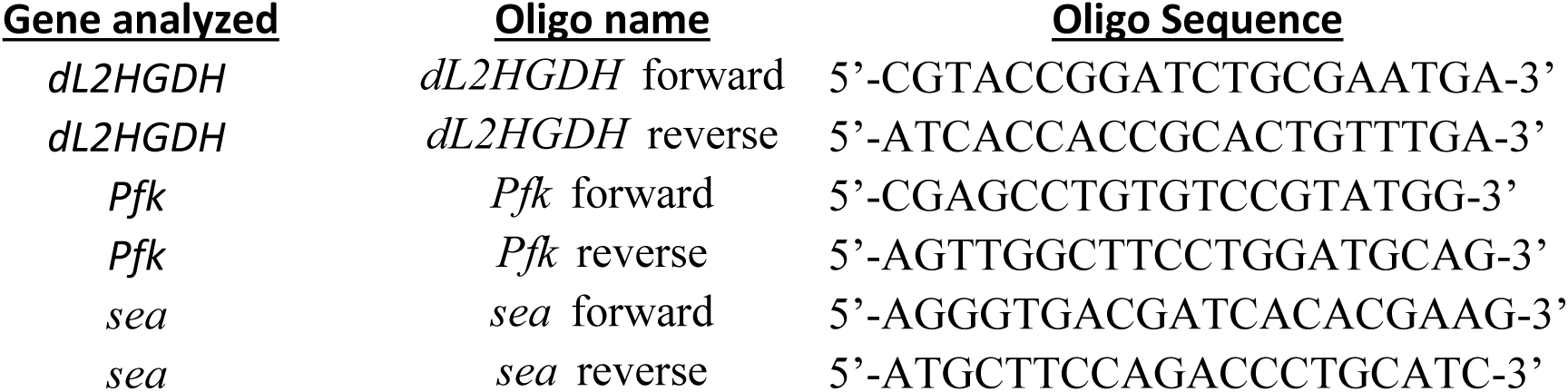
Oligos used for qRT-PCR analysis

## References

Anders S, Pyl PT, Huber W. 2015. HTSeq--a Python framework to work with high-throughput sequencing data. Bioinformatics 31: 166–169.

Backhaus B, Sulkowski E, Schlote F. 1984. A semi-synthetic, general-purpose medium for *Drosophila melanogaster*. Dros Inf Serv 60: 210–212.

Becker-Kettern J, Paczia N, Conrotte JF, Kay DP, Guignard C, Jung PP, Linster CL. 2016. *Saccharomyces cerevisiae* Forms D-2-Hydroxyglutarate and Couples Its Degradation to D-Lactate Formation via a Cytosolic Transhydrogenase. The Journal of biological chemistry 291: 6036–6058.

Burr SP, Costa AS, Grice GL, Timms RT, Lobb IT, Freisinger P, Dodd RB, Dougan G, Lehner PJ, Frezza C et al. 2016. Mitochondrial Protein Lipoylation and the 2-Oxoglutarate Dehydrogenase Complex Controls HIF1alpha Stability in Aerobic Conditions. Cell Metabolism 24: 740–752.

Carrisi C, Madeo M, Morciano P, Dolce V, Cenci G, Cappello AR, Mazzeo G, Iacopetta D, Capobianco L. 2008. Identification of the *Drosophila melanogaster* mitochondrial citrate carrier: Bacterial expression, reconstitution, functional characterization and developmental distribution. Journal of Biochemistry 144: 389–392.

Dolce V, Cappello AR, Capobianco L. 2014. Mitochondrial tricarboxylate and dicarboxylate-tricarboxylate carriers: from animals to plants. IUBMB Life 66: 462–471.

Fan J, Teng X, Liu L, Mattaini KR, Looper RE, Vander Heiden MG, Rabinowitz JD. 2015. Human phosphoglycerate dehydrogenase produces the oncometabolite D-2-hydroxyglutarate. ACS chemical biology 10: 510–516.

Intlekofer AM, Dematteo RG, Venneti S, Finley LW, Lu C, Judkins AR, Rustenburg AS, Grinaway PB, Chodera JD, Cross JR et al. 2015. Hypoxia Induces Production of L-2-Hydroxyglutarate. Cell Metabolism.

Intlekofer AM, Wang B, Liu H, Shah H, Carmona-Fontaine C, Rustenburg AS, Salah S, Gunner MR, Chodera JD, Cross JR et al. 2017. L-2-Hydroxyglutarate production arises from noncanonical enzyme function at acidic pH. Nature chemical biology 13: 494–500.

Jiang L, Boufersaoui A, Yang C, Ko B, Rakheja D, Guevara G, Hu Z, DeBerardinis RJ. 2017. Quantitative metabolic flux analysis reveals an unconventional pathway of fatty acid synthesis in cancer cells deficient for the mitochondrial citrate transport protein. Metabolic engineering 43: 198–207.

Kemp RG, Foe LG. 1983. Allosteric regulatory properties of muscle phosphofructokinase. Molecular and cellular biochemistry 57: 147–154.

Kim D, Langmead B, Salzberg SL. 2015. HISAT: a fast spliced aligner with low memory requirements. Nat Methods12: 357–360.

Kranendijk M, Struys EA, Salomons GS, Van der Knaap MS, Jakobs C. 2012. Progress in understanding 2-hydroxyglutaric acidurias. J Inherit Metab Dis 35: 571–587.

Li H, Chawla G, Hurlburt AJ, Sterrett MC, Zaslaver O, Cox J, Karty JA, Rosebrock AP, Caudy AA, Tennessen JM. 2017. *Drosophila* larvae synthesize the putative oncometabolite L-2-hydroxyglutarate during normal developmental growth. Proceedings of the National Academy of Sciences of the United States of America 114: 1353–1358.

Li H, Tennessen JM. 2017. Methods for studying the metabolic basis of Drosophila development. Wiley Interdiscip Rev Dev Biol 6: e280.

Lommen A. 2009. MetAlign: Interface-Driven, Versatile Metabolomics Tool for Hyphenated Full-Scan Mass Spectrometry Data Preprocessing. Analytical Chemistry 81: 3079–3086.

Losman JA, Kaelin WG, Jr. 2013. What a difference a hydroxyl makes: mutant IDH, (R)-2-hydroxyglutarate, and cancer. Genes & Dev 27: 836–852.

Love MI, Huber W, Anders S. 2014. Moderated estimation of fold change and dispersion for RNA-seq data with DESeq2. Genome Biol 15: 550.

Martin M. 2011. Cutadapt removes adapter sequences from high-throughput sequencing reads. EMBnetjournal 17: 10–12.

Morciano P, Carrisi C, Capobianco L, Mannini L, Burgio G, Cestra G, De Benedetto GE, Corona DF, Musio A, Cenci G. 2009. A conserved role for the mitochondrial citrate transporter Sea/SLC25A1 in the maintenance of chromosome integrity. Hum Mol Genet 18: 4180–4188.

Muhlhausen C, Salomons GS, Lukacs Z, Struys EA, van der Knaap MS, Ullrich K, Santer R. 2014. Combined D2-/L2-hydroxyglutaric aciduria (SLC25A1 deficiency): clinical course and effects of citrate treatment. J Inherit Metab Dis 37: 775–781.

Mullen AR, Wheaton WW, Jin ES, Chen PH, Sullivan LB, Cheng T, Yang Y, Linehan WM, Chandel NS, DeBerardinis RJ. 2011. Reductive carboxylation supports growth in tumour cells with defective mitochondria. Nature 481: 385–388.

Muntau AC, Röschinger W, Merkenschlager A, Van Der Knaap M, Jakobs C, Duran M, Hoffmann G, Roscher A. 2000. Combined D-2-and L-2-hydroxyglutaric aciduria with neonatal onset encephalopathy: a third biochemical variant of 2-hydroxyglutaric aciduria? Neuropediatrics 31: 137–140.

Nadtochiy SM, Schafer X, Fu D, Nehrke K, Munger J, Brookes PS. 2016. Acidic pH Is a Metabolic Switch for 2-Hydroxyglutarate Generation and Signaling. The Journal of biological chemistry 291: 20188–20197.

Nota B, Struys EA, Pop A, Jansen EE, Ojeda MRF, Kanhai WA, Kranendijk M, van Dooren SJM, Bevova MR, Sistermans EA et al. 2013. Deficiency in SLC25A1, Encoding the Mitochondrial Citrate Carrier, Causes Combined D-2-and L-2-Hydroxyglutaric Aciduria. American Journal of Human Genetics 92: 627–631.

Oldham WM, Clish CB, Yang Y, Loscalzo J. 2015. Hypoxia-Mediated Increases in l-2-hydroxyglutarate Coordinate the Metabolic Response to Reductive Stress. Cell Metab.

Palmieri F. 2004. The mitochondrial transporter family (SLC25): physiological and pathological implications. Pflugers Arch 447: 689–709.

Palmieri F. 2013. The mitochondrial transporter family SLC25: identification, properties and physiopathology. Molecular aspects of medicine 34: 465–484.

Pogson CI, Randle PJ. 1966. The control of rat-heart phosphofructokinase by citrate and other regulators. The Biochemical journal 100: 683–693.

Prasun P, Young S, Salomons G, Werneke A, Jiang YH, Struys E, Paige M, Avantaggiati ML, McDonald M. 2015. Expanding the Clinical Spectrum of Mitochondrial Citrate Carrier (SLC25A1) Deficiency: Facial Dysmorphism in Siblings with Epileptic Encephalopathy and Combined D,L-2-Hydroxyglutaric Aciduria. JIMD Rep 19: 111–115.

Reinecke CJ, Koekemoer G, van der Westhuizen FH, Louw R, Lindequie JZ, Mienie LJ, Smuts I. 2012. Metabolomics of urinary organic acids in respiratory chain deficiencies in children. Metabolomics 8: 264–283.

Rzem R, Veiga-da-Cunha M, Noel G, Goffette S, Nassogne MC, Tabarki B, Scholler C, Marquardt T, Vikkula M, Van Schaftingen E. 2004. A gene encoding a putative FAD-dependent L-2-hydroxyglutarate dehydrogenase is mutated in L-2-hydroxyglutaric aciduria. Proc Natl Acad Sci U S A 101: 16849–16854.

Struys EA, Salomons GS, Achouri Y, Van Schaftingen E, Grosso S, Craigen WJ, Verhoeven NM, Jakobs C. 2005a. Mutations in the D-2-hydroxyglutarate dehydrogenase gene cause D-2-hydroxyglutaric aciduria. Am J Hum Genet 76: 358–360.

Struys EA, Verhoeven NM, Ten Brink HJ, Wickenhagen WV, Gibson KM, Jakobs C. 2005b.) Kinetic characterization of human hydroxyacid-oxoacid transhydrogenase: relevance to D-2-hydroxyglutaric and gamma-hydroxybutyric acidurias. J Inherit Metab Dis 28: 921–930.

Teng X, Emmett MJ, Lazar MA, Goldberg E, Rabinowitz JD. 2016. Lactate Dehydrogenase C Produces S-2-Hydroxyglutarate in Mouse Testis. ACS chemical biology 11: 2420–2427.

Tornheim K, Lowenstein JM. 1976. Control of phosphofructokinase from rat skeletal muscle. Effects of fructose diphosphate, AMP, ATP, and citrate. The Journal of biological chemistry 251: 7322–7328.

Tyrakis PA, Palazon A, Macias D, Lee KL, Phan AT, Velica P, You J, Chia GS, Sim J, Doedens A et al. 2016. S-2-hydroxyglutarate regulates CD8+ T-lymphocyte fate. Nature.

Usenik A, Legiša M. 2010. Evolution of allosteric citrate binding sites on 6-phosphofructo-1-kinase. PLoS One 5: e15447.

Xia J, Wishart DS. 2016. Using MetaboAnalyst 3.0 for Comprehensive Metabolomics Data Analysis. Current protocols in bioinformatics 55: 14.10.11–14.10.91.

Ye D, Guan KL, Xiong Y. 2018. Metabolism, Activity, and Targeting of D- and L-2-Hydroxyglutarates. Trends Cancer 4: 151–165.

